# Population history of Swedish cattle breeds: estimates and model checking

**DOI:** 10.1101/2024.10.03.616479

**Authors:** Dolapo Adepoju, J Ingemar Ohlsson, Tomas Klingström, Elisenda Rius-Vilarrasa, Anna M Johansson, Martin Johnsson

## Abstract

In this work, we use linkage disequilibrium-based methods to estimate recent population history from genotype data in Swedish cattle breeds, as well as international Holstein and Jersey cattle data for comparison. Our results suggest that these breeds have been effectively large up until recently, when they declined around the onset of systematic breeding. The inferred trajectories were qualitatively similar, with a large historical population and one decline. We used population genetic simulation to check the inferences. When comparing simulations from the inferred population histories to real data, the proportion low-frequency variants in real data was different than was implied by the inferred population histories, and there was somewhat higher genomic inbreeding in real data than implied by the inferred histories. The inferred population histories imply that much of the variation we see today is transient, and it will be lost as the populations settle into a new equilibrium, even if efforts to maintain effective population size at current levels are successful.

## Introduction

Population history affects the amount and structure of genetic variation, and thus the efficacy of selection, the accuracy of genomic prediction, and the maintenance of diversity for the future. In population genetic terms, the history of a population can be described as a trajectory of effective population size over time. The effective population size is the size of an idealised population with an equivalent loss of genetic diversity or increase in inbreeding to the real population (reviewed by [1]).

Various methods exist for estimating population history from molecular or pedigree data. In this work, we use the linkage disequilibrium-based method GONE [2] to estimate recent population history from molecular data in Swedish cattle breeds, as well as publicly available international Holstein and Jersey cattle data for comparison. A rationale for concentrating on recent population history is that cattle breeds as we know them today were created relatively recently. Linkage disequilibrium-based methods perform well for recent history up to 100-200 generations ago [2, 3] and are relatively insensitive to selection [4], but can be biased by population structure, admixture and gene flow [5].

The main commercially important cattle breeds in Sweden are Holstein, Swedish Red and Jersey cattle. There are also eight local breeds (listed in FAO-DADIS [6]) that are under conservation efforts by breed societies, supported by the state. These breeds have all gradually decreased in census size as they have been replaced by international dairy and beef breeds [7–9], and according to the DAD-IS report, they are all considered at risk. On the other hand, the local breeds have not had the same intensity of selection as the commercial breeds. As a consequence, commercial breeds often also have small current effective population sizes [10–12]. Some of the local breeds have a known history of admixture. Swedish Red Cattle was formed in the late 19^th^ century with contributions from Shorthorn, Ayrshire and local Swedish cattle [13, 14]. Swedish Polled (SKB) are a result of merging the breed societies of Fjäll cattle and Swedish Red Polled in 1938 and then from the 1960s also some introgression from other breeds to improve milk production [13]; the samples included in this study are known to have a majority of Fjäll ancestry. The Fjällnära cattle originates from four genetically different herds and in our genomic data we have representatives from all four ancestries [15, 16].

In this paper, we estimate the recent population history of Swedish breeds, with international Holstein and Jersey cattle as comparison. We also use population genetic simulation to check the inferences that we have made. In the process, we provide simulation code for the inferred population histories that can be used for further research.

## Methods

### Data

For the Swedish cattle breeds, we used high-density SNP chip data generated by [16]. These animals were genotyped on a 150k Illumina array. For comparison, we extracted samples from Holstein and Jersey cattle from the public release of the 1000 Bull genomes project [17], run 9, downloaded from the European Nucleotide Archive (project accession PRJEB56689). These data, based on Illumina sequencing from many partners, are provided as genotypes in variant call format. We used the filter information provided to include only single nucleotide variants that passed variant quality score recalibration. We used Plink version 1.90b6.21 [18], bcftools version 1.9 [19] and GATK version 4.2.6.1 [20] for filtering and format conversion of genotype data. Table 1 shows the number of animals and data type for each breed.

**Table 1.** Sample sizes and type of data available for each breed.

| Breed | Description | Data type | Sample size |
| --- | --- | --- | --- |
| Bohus Polled | Native breed at risk | SNP chip | 7 |
| Fjäll | Native breed at risk | SNP chip | 23 |
| Fjällnära | Native breed at risk | SNP chip | 16 |
| Red Polled | Native breed at risk | SNP chip | 18 |
| Ringamåla | Native breed at risk | SNP chip | 13 |
| Swedish Polled (SKB) | Native breed at risk | SNP chip | 12 |
| Swedish Holstein-Friesian | Commercial breed,<br>Swedish sample | SNP chip | 24 |
| Swedish Red | Native breed, not at risk,<br>commercial | SNP chip | 25 |
| Väne | Native breed at risk | SNP chip | 9 |
| Holstein 1000 Bulls | Commercial breed,<br>international sample | Sequence | 146 |
| Jersey 1000 Bulls | Commercial breed,<br>international sample | Sequence | 77 |

### Population history inference

To infer recent population history over the last 200 generations, we used the GONE software version 29/08/2021 [2]. GONE uses an equation to predict the pairwise linkage disequilibrium for a given population history, and a genetic algorithm to fit a population history to the data. We applied GONE to SNP chip or sequence data from each population without allele frequency filtering. In cases where there are more than 50,000 variants on a chromosome, GONE randomly subsets the variants. In keeping with recommendations of the authors, we subset the inferred population history to the last 200 generations. We also summarised the population histories for each breed by finding the generation with the biggest change in effective population size from the previous generation, and the mean effective population size in generations before this decline.

We also tested the classical equation-based model for estimating population history from linkage disequilibrium [21–23], as implemented in SNeP version 1.11 [24], on the same data.

### Population genetic simulation

In order to check the validity of the population history inferences, we performed population genetic simulations. We checked the performance of GONE and SNeP on data simulated from known simplified histories. We also simulated data from population histories inferred by GONE and compared to real data with respect to statistics not directly used for inference (as recommended by [25]). Simulations were performed with msprime version 1.3.1 [26].

In order to avoid problems with coalescent simulation of small effective population sizes, which tend to underestimate long-range linkage disequilibrium in particular [27], we used a hybrid simulation strategy with coalescent simulation in the large historical population followed by discrete Wright—Fisher simulation for the last 200 generations, adjusting the population size at pre-defined times. Finally, we extracted a population sample from the last generation, and extracted genotypes at either a simulated 100k SNP chip or used all variants to simulate whole-genome sequence data.

### Simulations to check the performance of inference methods

We applied GONE and SNeP to data simulated from three different population histories, sampling 20 individuals from the last generation. We replicated each simulation 10 times, ran population history inference, and then plotted the inferred and true population history together.

We simulated these scenarios:

- The cattle population history estimated by Macleod et al. [28].
- A simple declining population history with a large historical population size of 5000 followed by a decline 50 generations ago to 100.
- Decline and recovery, with a large historical population size of 5000 followed by a decline 50 generations ago to 50, and an increase 20 generations ago to 100.

This gives us insight into the error profile of the methods, when applied to declining population histories similar to the ones we expect for recent livestock histories.

### Simulation checks of estimated population histories

We simulated data from the inferred population histories from Fjäll, Red Polled, Swedish Holstein-Friesian, Swedish Red, Holstein, Jersey and the published Holstein population history of Macleod et al. [28]. For comparison, we used the SNP chip data from Swedish breeds and the 1000 Bull genomes data for Holstein and Jersey, i.e., the same data that use for inference. We also included sequence data from 7 Fjäll cattle and 9 Swedish Red Polled cattle, previously published by [29].

We used the same number of simulated animals as in the empirical data. For comparison with SNP chip data, we subsampled SNPs randomly to a 100k SNP chip. For comparison with sequence data, we used all variants.

From both empirical and simulated data, we used Plink to estimate:

- Minor allele frequency spectrum calculated in frequency windows. We used windows of 0.1 for most breeds and of 0.05 for the 1000 Bull genomes Holstein and Jersey cattle, where the sample size is greater.
- Inbreeding coefficient based on homozygous genotypes *F*_*H*0*M*_.
- Inbreeding coefficient based on runs of homozygosity *F*_*R*0*H*_. Runs of homozygosity were detected with Plink (using default settings for SNP chip data, and allowing for 3 heterozygous and 10 missing markers per window for sequence data), meaning that this is inbreeding coefficient based on runs of at least 1 Mbp length. The inbreeding coefficient was calculated by dividing the total length of runs in an animal by the autosomal genome length.

We plotted simulation replicates against the estimates from real data. This gives us insight into how well the inferred population histories match real data, based on features not directly used in inference.

### Simulation of future genetic variation

Finally, to illustrate the consequences of the population histories for genetic variation, we ran simulations where each population history was extended with another 200 generations at the estimated final effective population size for a total length of 400 generations. For this simulation, we only included one chromosome, the length of cattle chromosome 1. Every 10 generations, we sampled 20 individuals and estimated the pairwise nucleotide diversity (*π*) within the sample, which we plotted to illustrate the decline in genetic diversity over time. The diversity calculation was performed with tskit version 0.5.6 [30].

## Data and code availability

The SNP chip data are available in the Dryad repository at https://doi.org/10.5061/dryad.wdbrv15j4. The 1000 Bull genomes data are available from the European Nucleotide archive at project accession PRJEB56689. The sequence data from Swedish cattle are available from the European Nucleotide archive at project accession PRJEB60564. The scripts for data analysis are available at https://www.github.com/mrtnj/cattle_population_history.

## Results

### Inferred population histories

The inferred population histories all showed a qualitatively similar pattern, with historical population size in the thousands, and a recent decline. Figure 1 shows the estimates for the Swedish breeds based on SNP chip data, showing the breeds with at least 10 animals sampled with the exception of Fjällnära cattle, which showed a pathological inferred history indicative of poor model fit. Figure 2 shows the estimates for Holstein and Jersey samples from the 1000 Bull genomes data, with comparison to previously published histories [28, 31].

**Figure 1.**
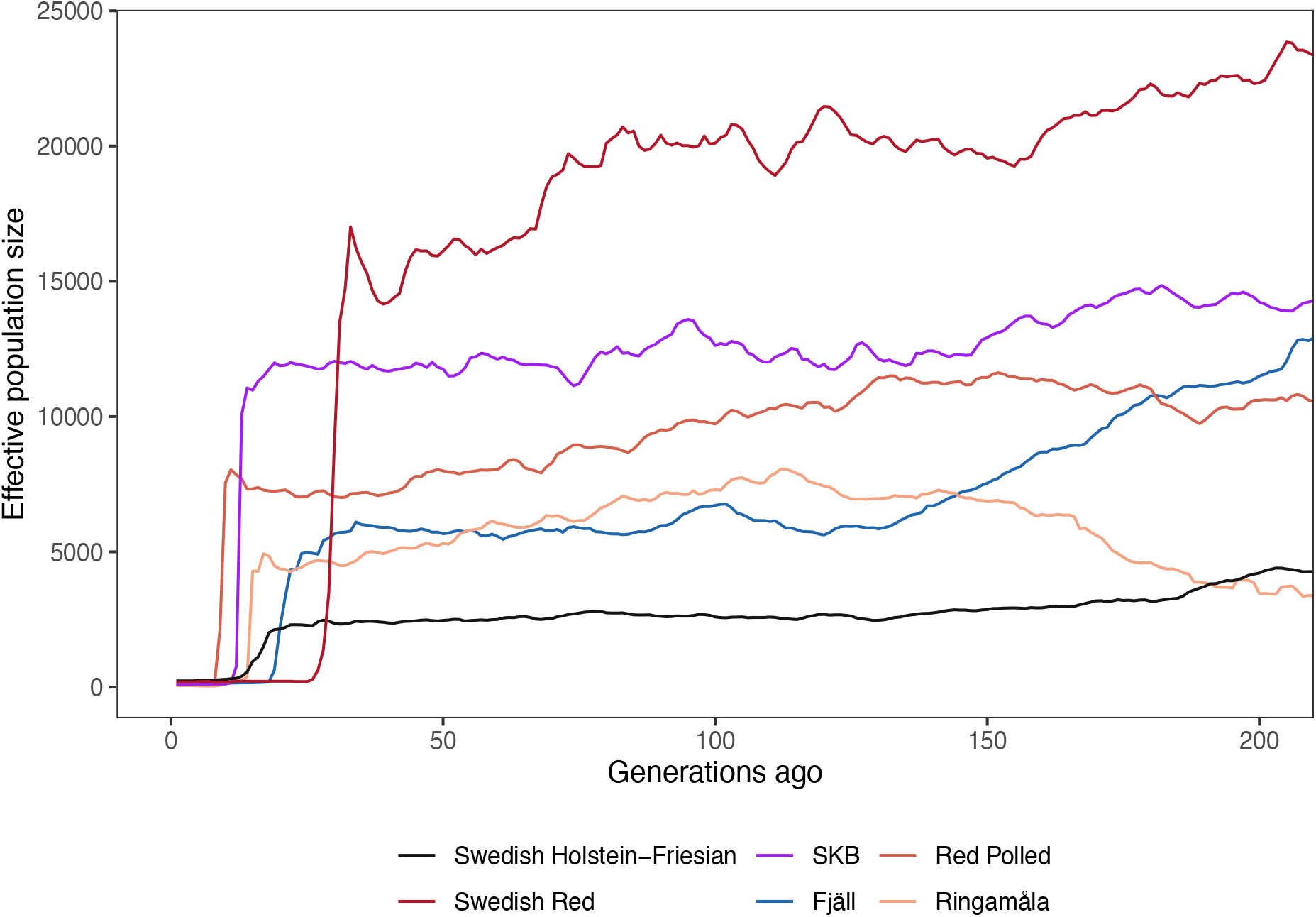
Population histories of Swedish cattle breeds, inferred by GONE from SNP chip data. The horizontal axis shows time in generations, running backwards. The vertical axis shows the estimated effective population size. The figure shows breeds with a sample size greater than 10, excluding the Fjällnära breed.

**Figure 2.**
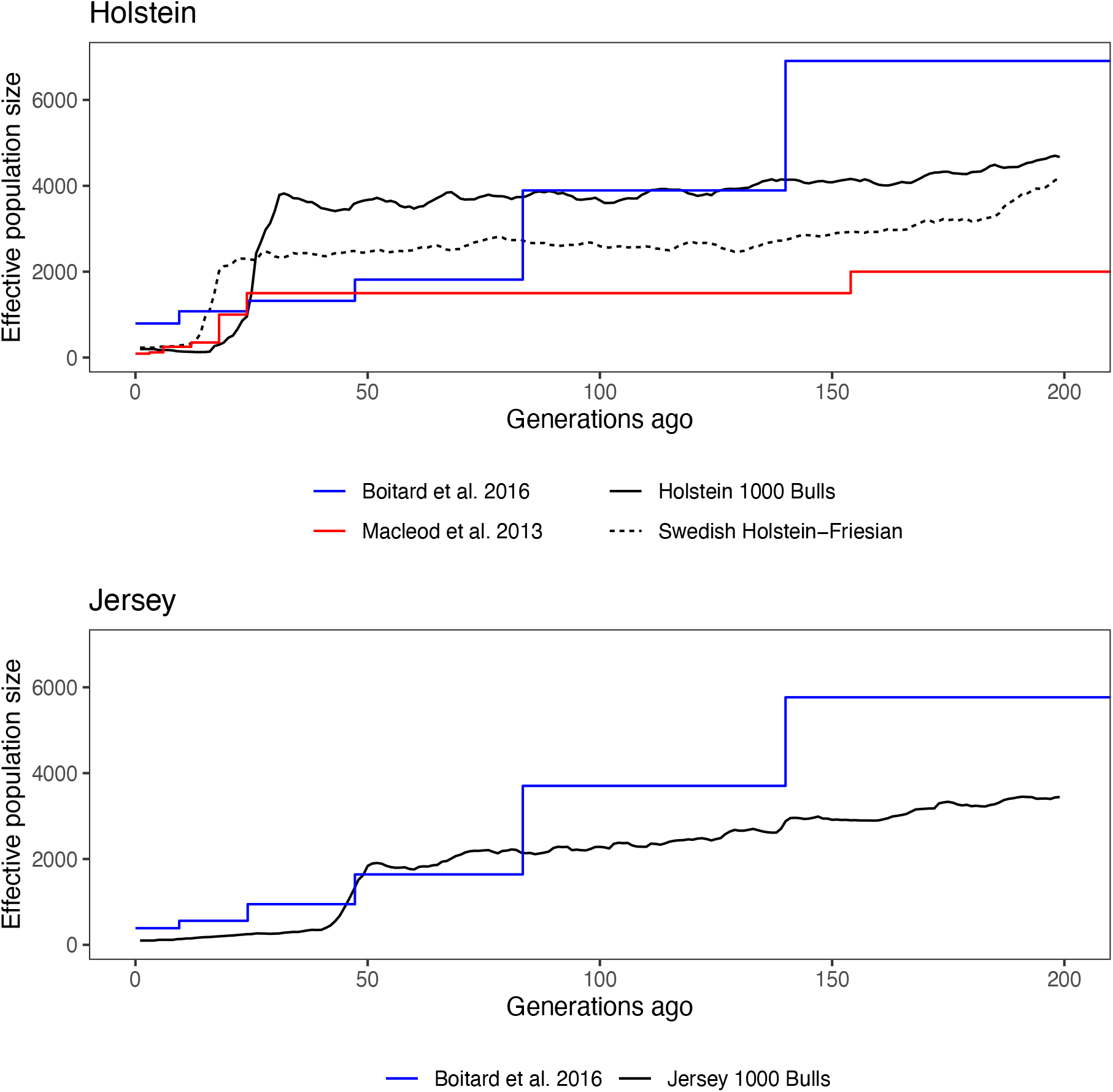
Population histories of international Holstein and Jersey cattle, inferred by GONE from 1000 Bull genomes genotypes; the dashed line shows Swedish Holstein—Friesian for comparison. The horizontal axis shows time in generations, running backwards. The vertical axis shows the estimated effective population size. The red and blue lines show published population histories.

Supplementary Figure 1 shows the inferred history for every breed on its own scale. Supplementary Table 1 shows the approximate timing of the decline, the mean effective population size before the decline. All inferred histories are provided in Supplementary Dataset The estimated current effective population sizes of all breeds ranged from around 20 to around 200. Figure 3 shows the estimated current effective population sizes of all breeds. The numerically small traditional breeds, such as Väne at an effective population size of 20, Fjällnära at 24 and Bohus Polled at 39, tended to be smaller than the commercial breeds, with Swedish Red at 189, and Swedish Holstein-Friesian at 229. However, Fjäll cattle was estimated to have higher effective population size than Jersey, with 132 compared to 101.

**Figure 3.**
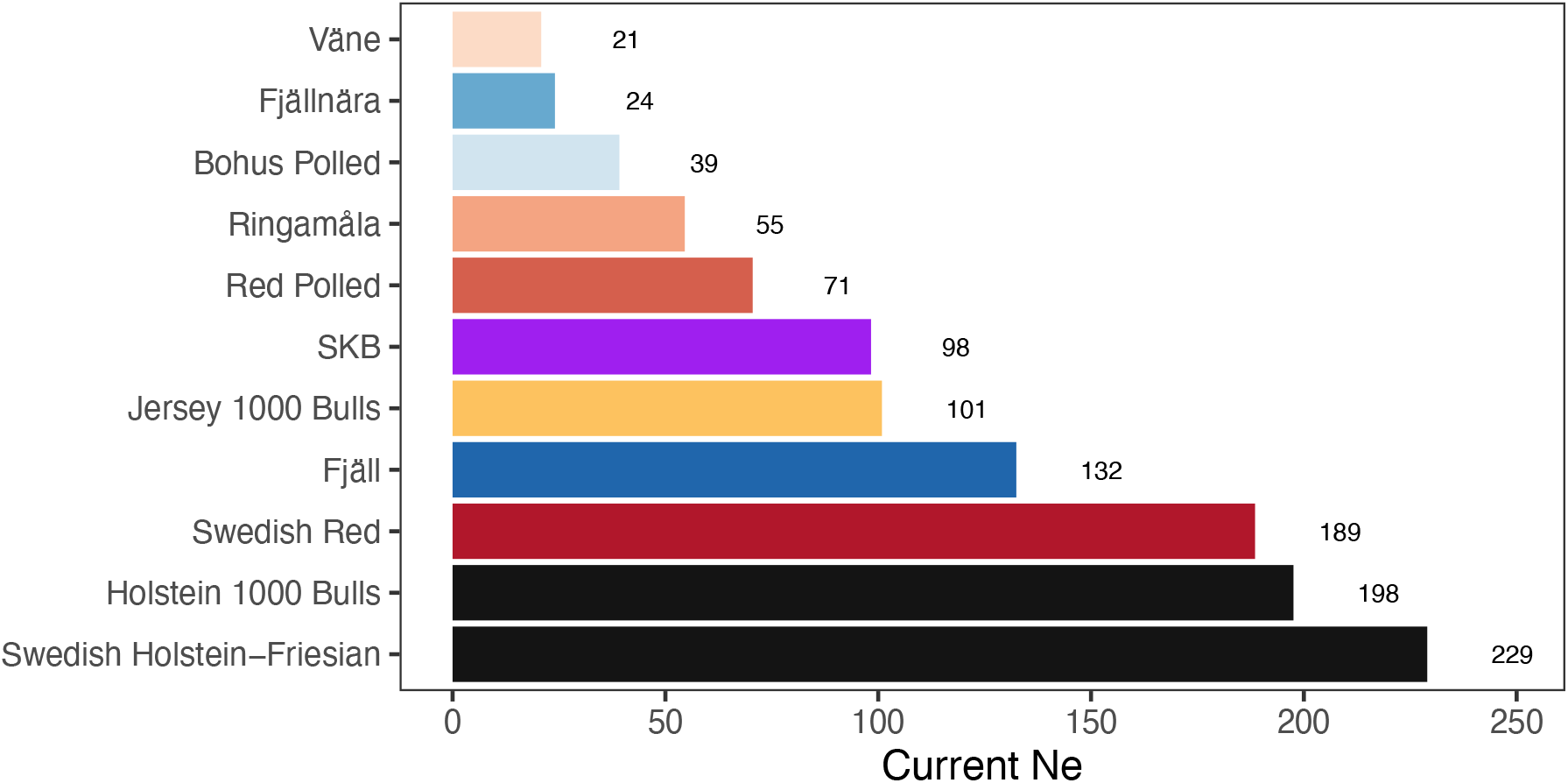
Estimates of current effective population size from GONE.

### Simulations

Inference on data simulated from known population histories suggested that the method was able to infer the shape of population histories with recent declines, but that there was substantial uncertainty in historical population size. Figure 4 shows estimated and simulated real population histories together. The mean absolute deviation from the true simulated effective population size was around 1000 from 50-100 generations ago. There was a systematic bias in estimates from the simple decline history, where in earlier generations (particularly pronounced before generation 150), the effective population size appeared to be spuriously growing. In the case of a recent extreme bottleneck with recovery, the method was unable to detect the previous population decline before the bottleneck, and instead inferred a spuriously small historical effective population size. Inference using SNeP performed poorly in simulation; results are shown in Supplementary Figure 2.

**Figure 4.**
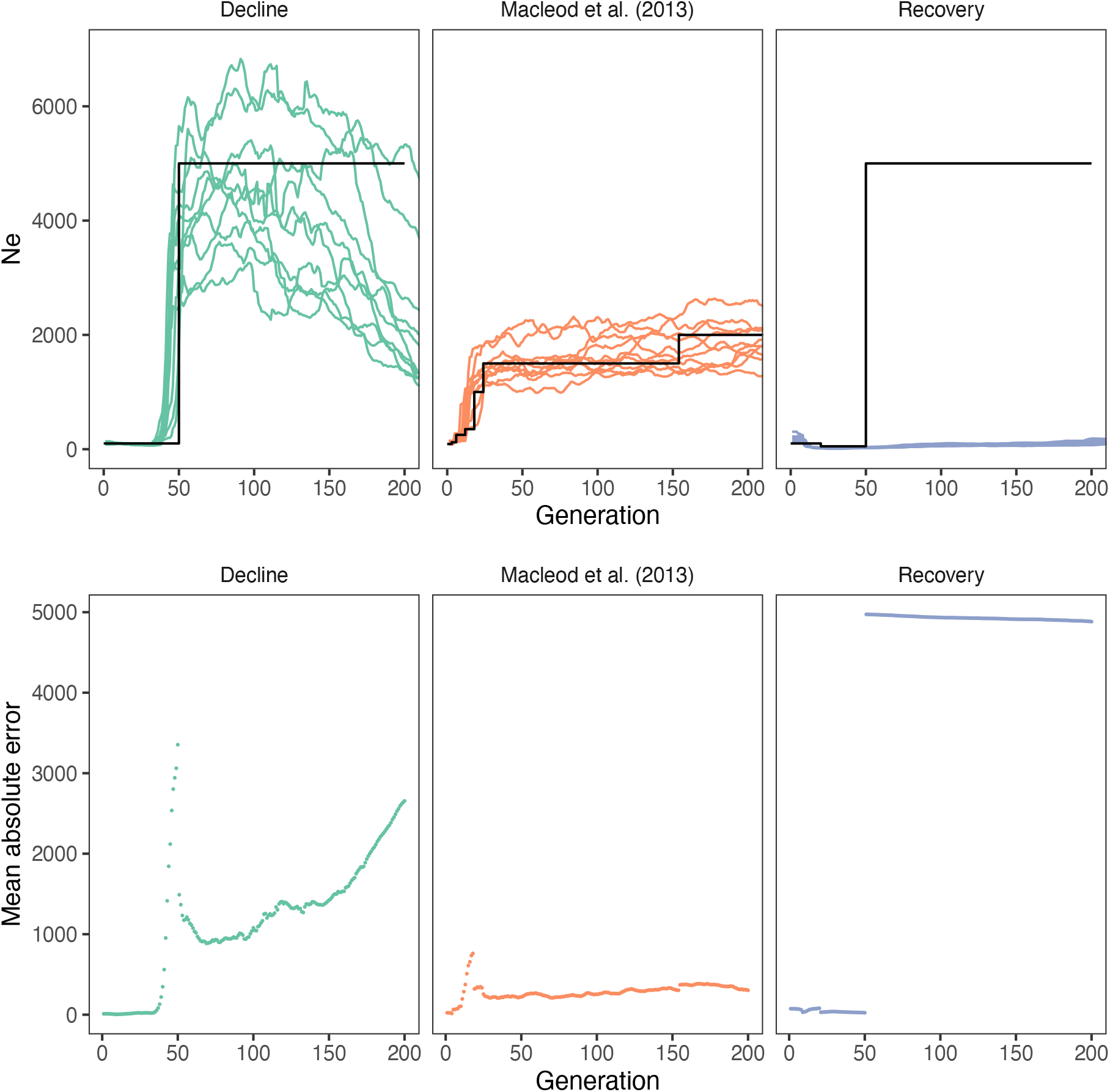
Population history inference with GONE applied to simulated fake data. The black lines show the true simulated population histories. The coloured lines show inferred population histories from replicated simulations. The points in the lower panel show the mean absolute deviation between real and inferred effective population size in each generation.

Simulations from inferred histories showed discrepancies between the allele frequency distribution implied by the inferred histories and the ones observed in real data. Figure 5 shows minor allele frequency distributions for data simulated from inferred population histories and estimated from real genotype data. In the SNP chip data from the Swedish cattle breeds, the simulated data showed an overrepresentation of rare variants compared to the real data. In sequence data from Fjäll and Red Polled cattle, there was an overrepresentation still, but smaller. In the sequence data from the 1000 Bulls, where sample sizes were bigger, there was an underrepresentation of rare variants in the simulated data compared to real sequence data. Simulations from the population history of Macleod et al. [28] also showed an underrepresentation of rare variants compared to real data.

**Figure 5.**
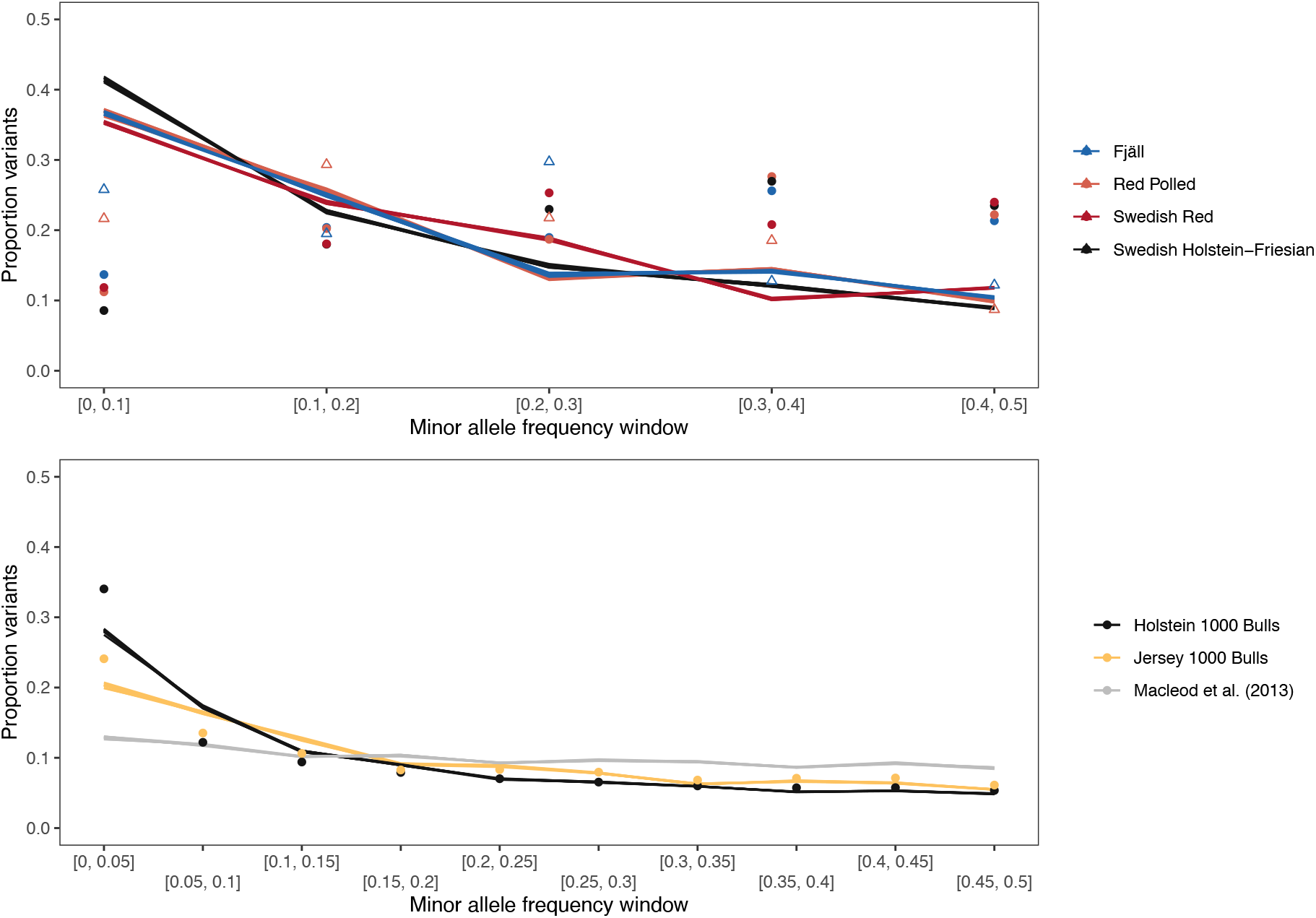
Comparison between the allele frequency distribution implied by inferred histories and real data. The lines show replicated simulations from the inferred population histories. Colours correspond to breeds. The round points show estimates from the data used for inference. In the case of the top panel, this is SNP chip data, and in the bottom panel 1000 Bull genomes genotypes. In the top panel, the open triangles show estimates from whole-genome sequence data from Fjäll and Red Polled cattle.

Simulation from inferred histories also suggested that these histories somewhat underestimate the level of inbreeding compared to real data. Figure 6 shows genomic inbreeding coefficients, as estimated by homozygosity or by runs of homozygosity, for data simulated from inferred population histories and estimated from real genotype data. With the exception of Fjäll cattle, the inferred population histories led to inbreeding coefficients that were on average lower than those observed in real data. The same was true for the population history of Macleod et al. [28] compared to Holstein data from the 1000 Bull genomes project. In particular, there were real animals with very high inbreeding coefficients, that were not generated by simulated data. In most cases, the mean of the simulated inbreeding coefficient was within a standard deviation of the real value.

**Figure 6.**
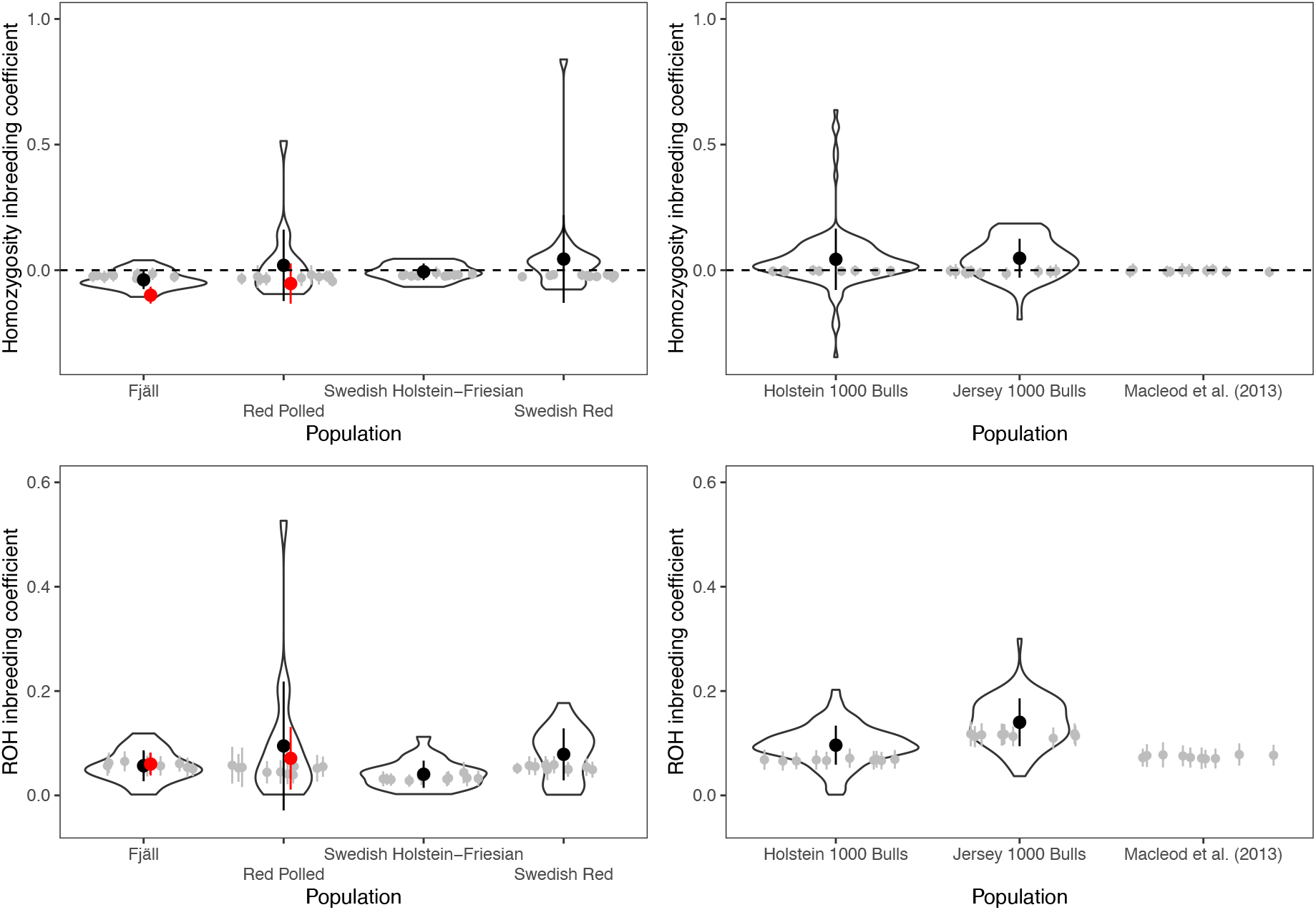
Comparison between the genomic inbreeding implied by inferred histories and real data. The violin plots show estimates from the data used for inference. The points and error bars show means and standard deviations of the genomic inbreeding estimated from simulated data. For the Swedish cattle, where inference is based on SNP chip data, the red points and bars show estimates from whole-genome sequence data from Fjäll and Red Polled cattle.

The inferred population histories imply populations that are not in equilibrium, and where there is transient genetic variation that will be lost over a long time. Figure 7 shows simulated trajectories of nucleotide diversity from four of the inferred population histories, extended by 200 generations where the population remains at the estimated current effective population size. The simulations showed nucleotide diversity remaining relatively constant up until the population size decline, and a steady decrease in diversity in the future, where the populations would not reach equilibrium within 200 generations.

**Figure 7.**
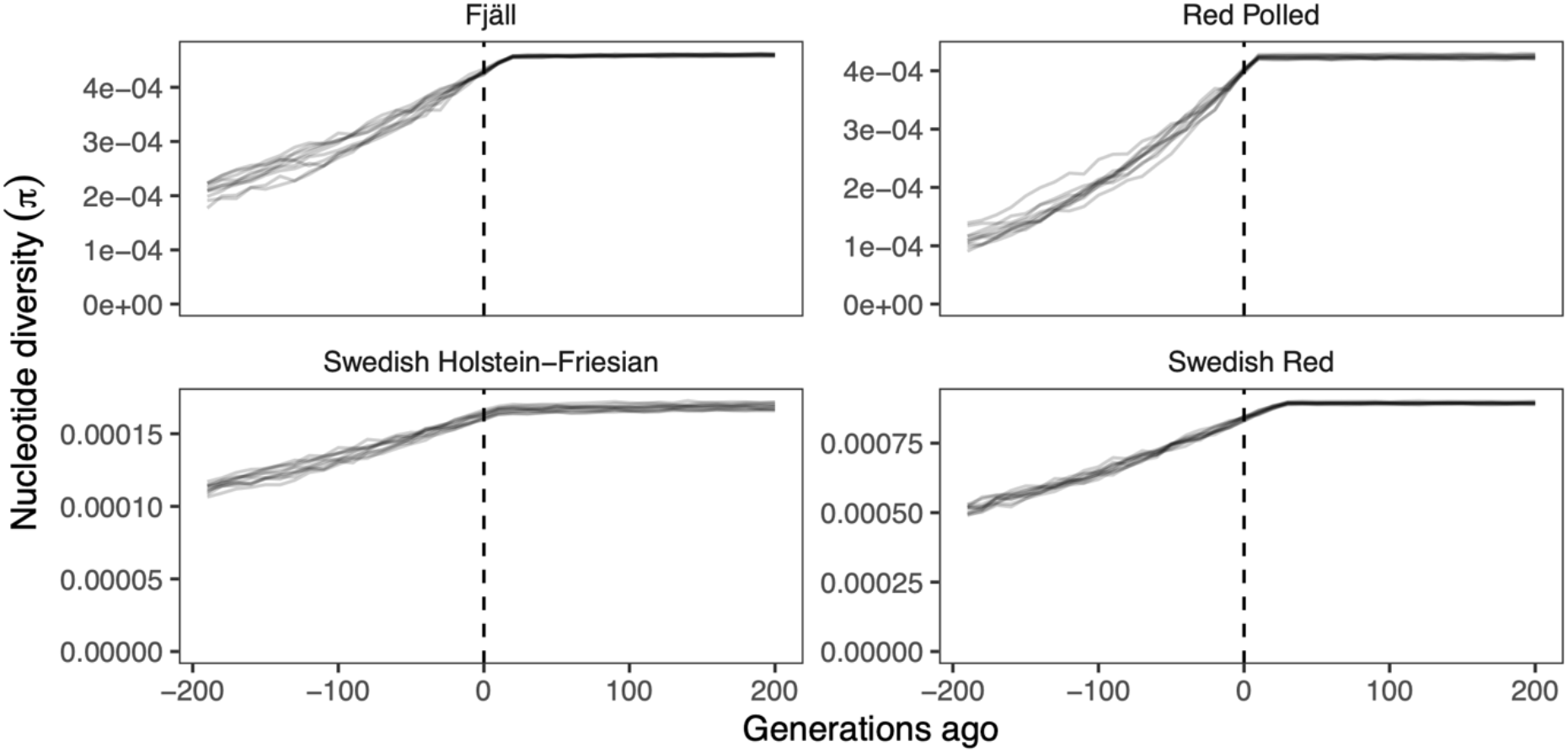
Future genetic diversity trajectories implied by inferred population histories. The lines show estimates of nucleotide diversity from samples of 20 individuals taken every 10 generations. For consistency with the other graphs, time is shown running backwards along the horizontal axis, with zero as the current time. Thus, the negative generation numbers represent future generations.

## Discussion

In this work, we estimated the recent population history of Swedish cattle breeds, with international Holstein and Jersey cattle as comparison. Our results suggest that these breeds have been effectively large up until recently, when they declined around the onset of systematic breeding. Simulations suggest that the inference method performs fairly well for cattle-like population histories with recent declines, but that it may perform poorly in populations with extreme bottlenecks and a recent recovery. When comparing simulations from the inferred population histories to real data, there were some discrepancies. There was a higher proportion low-frequency variants in 1000 Bull genomes data than is implied by the inferred population histories, and somewhat higher genomic inbreeding in real data than implied by the inferred histories.

We will discuss potential biases that affect these results, correspondence to known events, the relevance to quantitative genetic simulation of cattle breeding, and the implications for cattle genetic resources.

### Sources of error and bias

There are several sources of uncertainty and bias in effective population size estimation that should be kept in mind when interpreting our results.

The largest inferred historical effective population sizes in our analysis come from breeds that have a history of admixture and population structure. This is a known bias in the method [5], which assumes a sample from a closed and well-mixed population. Thus, the implausibly high historical effective population sizes of Swedish Red Cattle (around 20,000), Swedish Polled (SKB) (around 13,000) and Red Polled (around 10,000) should likely not be taken at face value. Historically, Swedish Red cattle was formed by crossing imported Shorthorn and Ayrshire with local Swedish cattle in the 18^th^ century [13, 14]. This scenario is similar to the creation of a synthetic breed simulated by Novo et al. to evaluate GONE’s performance. Their results suggest that the estimated timing of admixture, i.e., the inferred decline, is accurate but that the historical effective population size is overestimated compared to the combined size of the founding population [5]. More recently, the Swedish Red breed shares a genetic evaluation and semen providers with Danish and Finnish Red. Swedish Polled, on the other hand, is a product of administrative concerns, as the two breed societies for Red Polled (SKB) and Fjäll were joined in 1938 [13]. Admixture between Fjäll-derived and Red Polled-derived SKB cattle were probably not such a big factor, as the breeds are quite distinct both in terms of ancestry and phenotype. However, there was conscious crossing with other breeds, including Swedish Holstein—Friesian, Jersey and Finnish cattle [32]. Thus, both population structure and admixture likely contribute to inflating the inferred historical effective population size. The Red Polled breed shows historical population structure, with multiple ancestries including important influence from Norwegian and Finnish Red Polled cattle [13].

Fjällnära cattle showed a pathological inferred population history, with an exceptionally large recent population history and a steep decline. This pattern is similar to what the creators of GONE have shown to happen when there is strong population structure within the sample, which breaks the assumptions of the method [5]. Indeed, the Fjällnära breed is known to have strong population structure, with four distinct subpopulations [15].

Applying the inference method to simulated data suggests that there is substantial uncertainty in the magnitude of historical effective population size. This pattern is more pronounced in the simple decline scenario, which has a higher historical population size, compared to simulations of the Macleod et al. [28] population history. To fully explore this error and its scaling with population size would require more extensive simulations with ranges of parameters. Suffice to say that one should treat the estimated historical population sizes with caution. In simulations of declining populations, we also saw a bias in the early part of the population history where the effective population size appears to be growing when it was in fact simulated to be constant. This suggests that similar patterns for example in the inferred population history of Ringamåla cattle may not be trustworthy. In general, the error is higher for earlier generations, which should be expected.

Furthermore, all the Swedish cattle samples used in this study are non-random samples of their respective breed. We should expect, in general, an upward bias in historical effective population size. Similarly, the 1000 Bull genomes data combine samples from many different partners all over the world. This may explain why the estimated current effective population size, around 200, appears high compared to Holstein estimates from breeding programs [10, 12].

The simulated scenario of a recent bottleneck with recovery suggests that the method may struggle with inferring the population history after an extreme bottleneck. In the bottleneck scenario we simulated, the inferred population history detected a recent increase from a long-time low historical effective population size, suggesting that such an extreme bottleneck effectively erases information about previous historical population size. In the inferred histories, only Bohus Polled cattle, which is indeed a very small breed that has likely been subject to an extreme recent bottleneck, showed a pattern similar to this. However, the Bohus Polled history is also based on a small sample size, and shows other pathological features like a recent an instantaneous increase in effective population size. As a corollary, these results suggest that the bottlenecks in the other Swedish breeds have not been as extreme as the simulated case. However, we only simulated one parameter combination, and therefore caution is warranted.

Simulation-based checks of inferred population histories show discrepancies between the real allele frequency distributions and those implied by simulations. For the Swedish cattle data, there was an overrepresentation of low-frequency variants implied by the inferred history.

However, in the case of SNP chip data, this discrepancy may be due to ascertainment bias, where SNP chips are designed to be biased towards common variants. Further, the sequence data from Fjäll and Red Polled cattle come from small enough sample sizes (seven and nine cattle respectively) that they cannot estimate low frequencies well. In the case of the 1000 Bull genomes data, where sample sizes are greater, the inferred histories imply a smaller proportion of low-frequency variants than observed in real data.

Finally, our simulations suggest that the classical method implemented in SNeP do not perform well for population histories with recent declines. Problems with this method were pointed out already by Corbin et al. [23], but seemingly, the approach is still occasionally used in animal genetics.

### Correspondence to known events

The approximate time of the declines occur around ten to twenty generations ago, which corresponds to the mid to early 20^th^ century, assuming a generation time of around five years and that the cattle sampled were born in the late 1990s to early 2000s. This roughly corresponds to the onset of systematic breeding or the later decline due to the replacement of traditional breeds by international breeds. For example, Fjäll cattle the inferred decline occurs 19 generations or approximately 100 years ago (assuming a generation time of five years), and a breed standard was established in 1893 [13]. For Red Polled, the inferred decline occurs 9 generations or approximately 50 years ago; in the mid 20^th^ century the breed declined down to almost going extinct in the 1970s. For Swedish Red Cattle, the inferred decline occurs 29 generations or approximately 150 years ago. The original two breed societies for what would become Swedish Red were formed in 1892 and 1898. They were both already working with animals that were mixed between Shorthorn, Ayrshire and local Swedish cattle [14], suggesting that admixture had happened earlier. One exception is Jersey cattle, where the inferred decline occurs 47 generations or approximately 200 years ago. This is consistent with the breed’s island origin, where the population has been closed with no importation allowed from 1789 and a breed standard was established in 1834 [33].

In the case of Holstein and Jersey cattle, we can compare our inferred population histories to published histories [28, 31]. In both cases, the previously published histories go further back in time than the 200 generations used for our estimates, going back to 33,154 and 130,000 generations ago, respectively. Compared to previous histories, both our Swedish and international Holstein estimates suggest a higher population history more recently. The decline in population history is more dramatic compared to Macleod et al.’s inferred history, but less dramatic than Boitard et al.’s history, where the latter results in an unexpectedly high current effective population size close to 1000.

As mentioned above, our estimated current effective population sizes for Holstein appear high compared to current estimates from breeding programs (189 and 229 vs less than 50 in several breeding programs [10, 12]). Our estimate for Swedish Red cattle is lower (189 vs 226) compared to the pedigree-based analysis of Nyman et al. [11]. The estimate for Swedish Polled (SKB) was also lower than their pedigree-based estimate (98 vs 166). The pedigree-based estimates pertain to the whole pedigree based on data from 1960 to 2018, and in our inferred population histories we infer little change during this period. Thus, the estimates should have a comparable time frame.

The historical generation time of cattle is uncertain, as it changes with breeding practices. Above, we used a generation interval of 5 years, which is consistent with estimates from Red, Holstein and Jersey cattle before genomic prediction [11, 12]. After the introduction of genomic selection around 2010, the generation times of commercial cattle breeds have dramatically declined [12, 34]. The Swedish samples used in this study are not affected by this change, as the Swedish Red and Holstein—Friesian cattle were born before genomics, and the small breeds do not have genomic breeding. However, when viewing the 1000 Bull genomes estimates, one should keep in mind that the most recent generations likely have shorter generation intervals. This also affects the predictions of future decline in genetic variation. Commercial breeds like Holstein cattle have shorter generation times than a local breed under conservation breeding, with correspondingly higher rate of loss of variation over time.

Both Fjäll cattle and Red Polled cattle have suffered through documented recent genetic crises in the latter half of the 20^th^ century. The breeds declined severely, and efforts to rescue them were undertaken starting in the 1980s and 1990s [13, 32]. These efforts include collection and use of cryopreserved semen from old bulls as well as attention to ancestry and diversity rather than selection for trait improvement (as expressed in breeding plans [35–38]). The estimates of recent effective population sizes, which are comparable to estimates for international breeds, confirm that these efforts have to some extent been successful. For example, for Fjäll cattle, we estimated a current effective population size of 132 that appeared not to be declining further. We would like to emphasise that an inferred stable effective population sizes after the decline does not mean that there is no concern. These breeds are considered at risk, have decreasing census sizes, and serious concerns with economic viability for farmers.

At the same time, we note that our inferred population histories for these breeds do not show multiple declines, bottlenecks and recoveries. Instead, the inferred population histories simply appear like simple declines. This suggests that the method is unable to reconstruct complicated recent population history at sufficient resolution. In the breeds that suffered crises, it may be that the rescue efforts themselves are part of obscuring the traces of this history in the molecular data.

Also in Swedish Holstein and Red Cattle, genotyping of old bulls has shown that there has been fluctuations in inbreeding rate over the latter half of the 20^th^ century, where inbreeding decreased from the 1960s to 1980s and later increased again in Holstein but remained relatively stable in Red cattle [39]. That this signal can be recovered in historical genotypes suggests that it would be possible to estimate a more fine-grained population history from time series data.

### The relevance to quantitative genetic simulation

Our simulation-based checks of inferred population histories suggest a higher proportion of rare genetic variants than implied by previous estimates [28], and furthermore, that our inferred histories from 1000 Bull genomes data still lead to an underrepresentation of rare variants compared to the real genotypes. This has implications for the genetic architecture of complex traits in simulation, suggesting that simulated allele frequency distributions are often too uniform.

Such differences may not matter much to simulations that simply aim to predict the short-term response to selection in different breeding schemes. To a first approximation, response to selection depends on genetic variance, not details of genetic architecture. However, such properties may matter when simulations aim to describe mechanisms of responses to selection, genomic prediction accuracy and evolution of genetic variance, which may depend on linkage disequilibrium and genetic architecture [40–43]. As an example, Macleod et al. [44] found that when simulating genomic prediction with whole-genome sequence data, simulating a declining population history rather than a constantly large population destroyed the benefit of whole-genome sequence data.

With recent developments in population genetic simulation, using a more realistic population history when relevant is not particularly difficult. Tools like the community-maintained library of population histories stdpopsim [45, 46] will likely make it easier in the future. However, when simulating more realistic population histories, especially when attempting to simulate whole genomes or in combination with complications such as selection, computational constraints become a real problem. Our simulations comparing inferred population histories to real data suggest that is possible to create simulated data with similar or better agreement with reality—at least with respect to the allele frequency distribution and genomic inbreeding—than previous estimates without simulating ancient population history. This may save some time and computational resources.

### Implications for genetic resources

Due to a lack of resolution for recent events like multiple declines and bottlenecks, combined with substantial uncertainty, this type of molecular population history inference appears to have limited use in applied conservation of local breeds. For tracking generation-to-generation effective population size, one would need other methods that could be based on pedigree registration or on molecular monitoring.

It should be uncontroversial that genetic diversity of farm animals is decreasing. Our results suggest that, for these breeds, a large decrease in effective population size has happened over the last century, after the onset of systematic breeding. Effective population size has dramatically decreased both in numerically small breeds under conservation efforts and in international commercially used cattle breeds, and domestic cattle are now fragmented into effectively small populations. Even if current breeding programs succeed in limiting the future decrease in effective population size and maintain the current size, genetic variation will continue to decline for a long time. Much of the variation we see today is transient, and it will be lost as the populations settle into a new equilibrium. This, however, will take hundreds of generations.

It is an open question to what extent this loss of variation will impede response to selection or viability. Rules of thumb for viable effective population size are often cited as recommendations in conservation genetics (reviewed by [47, 48]). Short-term recommendations of effective population size above 50 or 100 are based on assumptions about tolerable levels of inbreeding depression over the next few generations. The smaller breeds in this study are below these thresholds. Long-term recommendations of effective population size above 500 or 1000 generations are based the equilibrium between mutational variance and loss of additive genetic variance in an infinitesimal model. On the other hand, Hill argued, using similar quantitative genetic arguments, that the supply of mutation in a population of effective size 100 may be sufficient for a sustained response to directional selection [49]. This depends on parameters that are hard to know, namely the mutational variance of quantitative traits, not just molecular mutation rates; our simulations, in contrast, only include neutral evolution. The main difference between Hill and the conservation geneticists is that Hill assumes a higher mutational variance. Current effective population sizes of these cattle breeds are mostly close to or below 100, and definitely below 500.

Looking to commercial breeds where there is data on quantitative traits, dairy cattle are not yet facing inbreeding depression that exceeds genetic gain [50]. There some signs of loss of genetic variance for breeding goal traits in pigs and Simmental cattle [51, 52], but not in a study of Holstein dairy cattle [53] and a chicken breeding program [54].

---

**Supplementary Figure 1.**
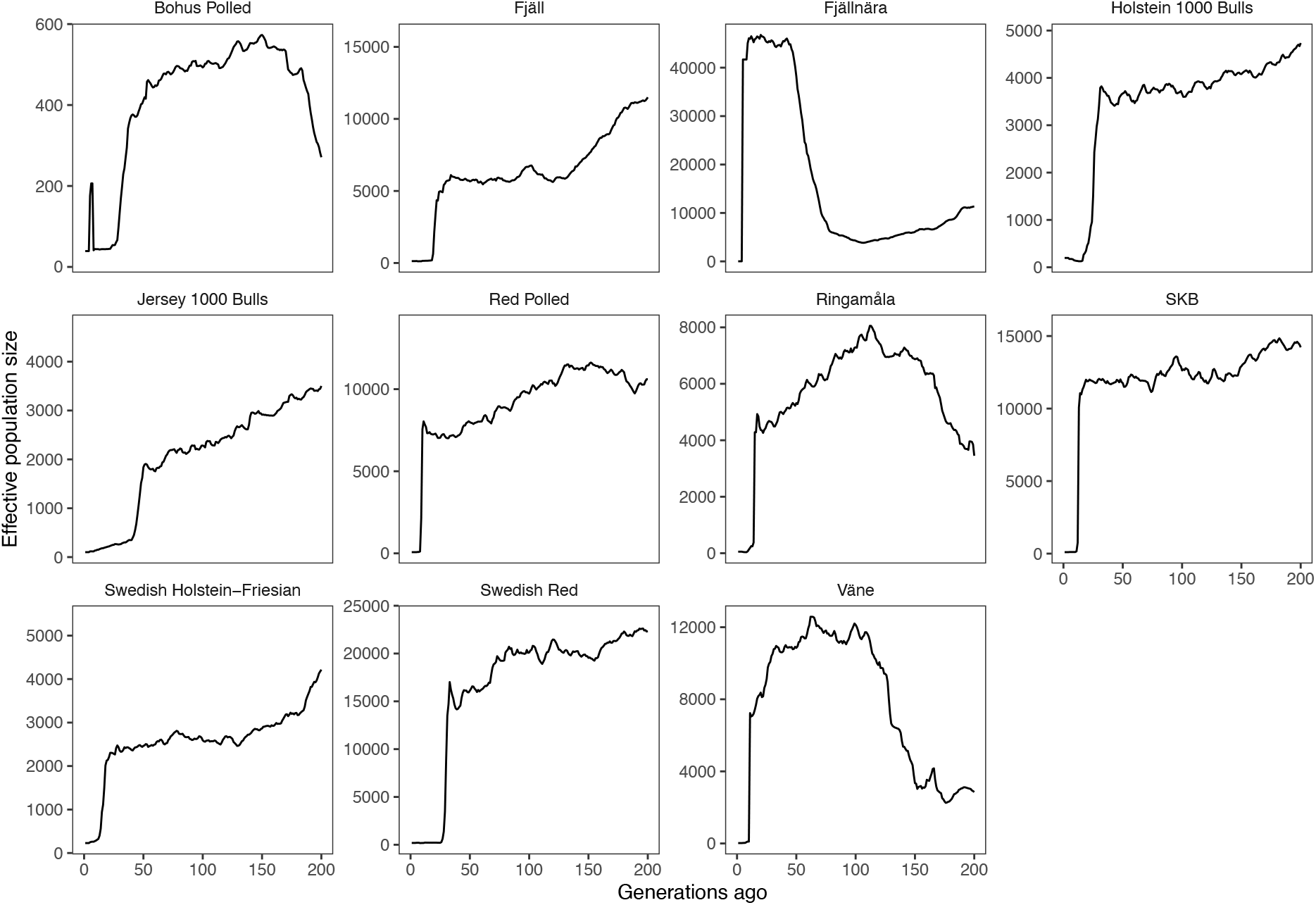
Population histories of inferred by GONE. The horizontal axis shows time in generations, running backwards. The vertical axis shows the estimated effective population size. This figure includes all estimated trajectories on their own scale.

**Supplementary Figure 2.**
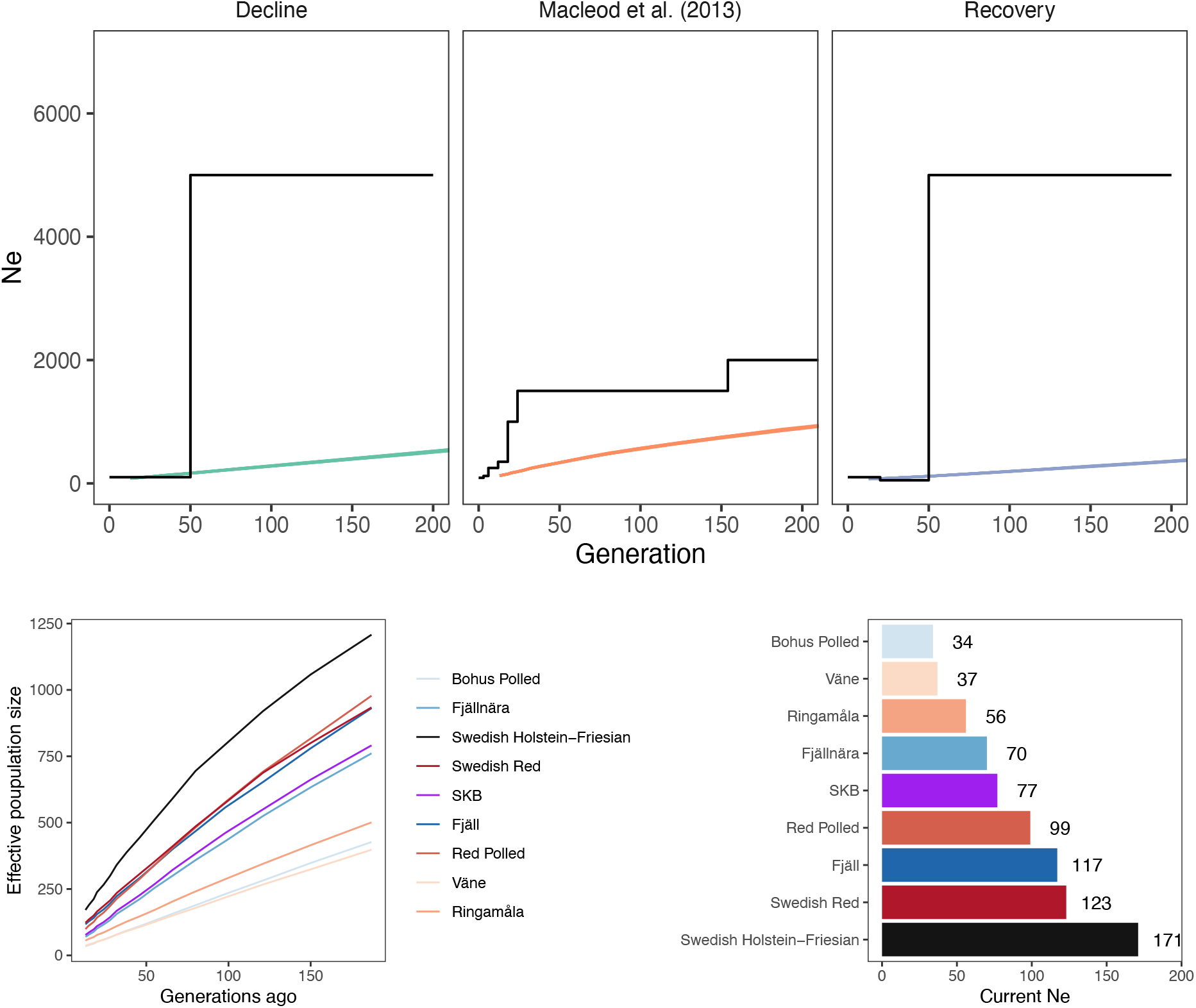
SNeP simulation and inference. The top panel shows the true simulated population histories in black and the inferred population histories from replicated simulations in colour. The bottom panels show population histories of Swedish cattle breeds, inferred by SNeP from SNP chip data, and the final estimated effective population size from SNeP.

**Supplementary Table 1.**
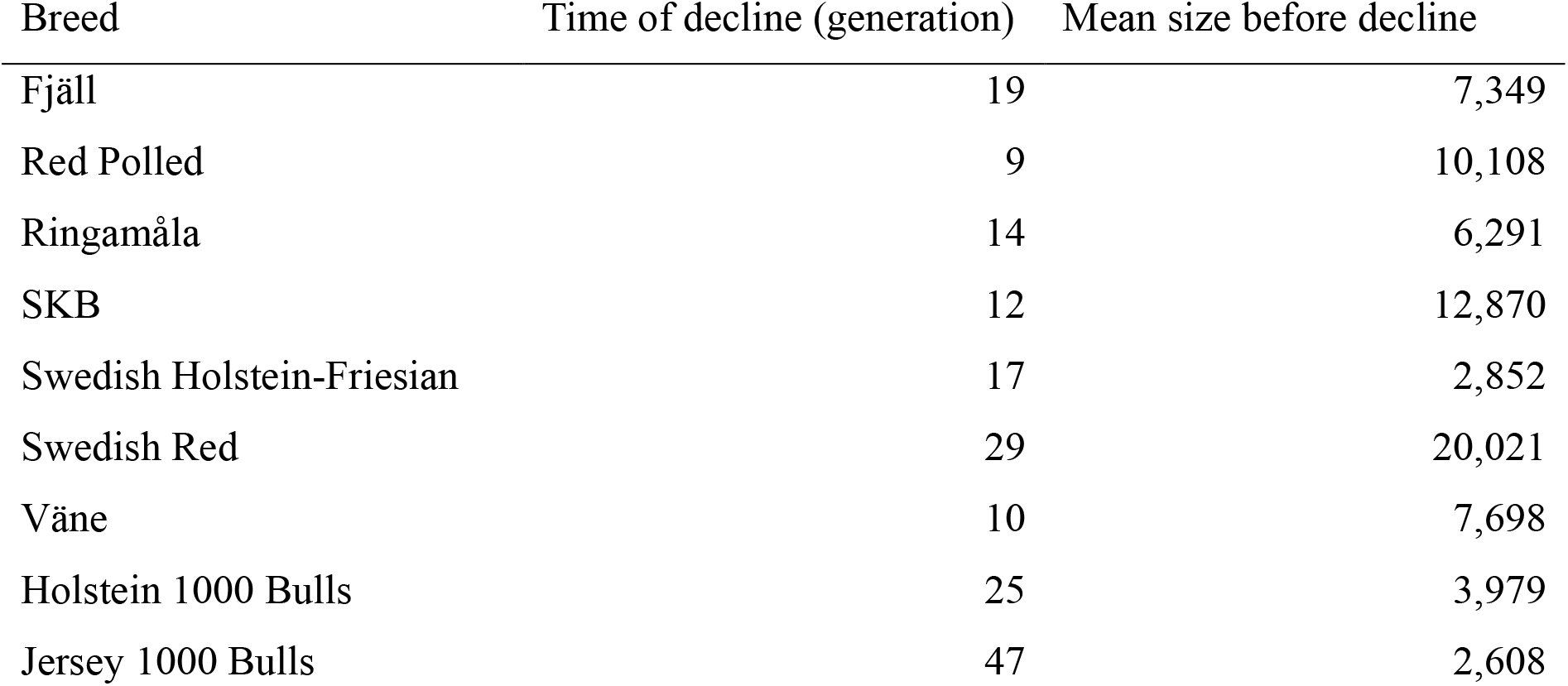
Time of the decline, estimated as the generation with the biggest decline in effective population size from the previous generation, and the mean effective population size before the decline, from population histories inferred by GONE.

| Breed | Time of decline (generation) | Mean size before decline |
| --- | --- | --- |
| Fjäll | 19 | 7,349 |
| Red Polled | 9 | 10,108 |
| Ringamåla | 14 | 6,291 |
| SKB | 12 | 12,870 |
| Swedish Holstein-Friesian | 17 | 2,852 |
| Swedish Red | 29 | 20,021 |
| Väne | 10 | 7,698 |
| Holstein 1000 Bulls | 25 | 3,979 |
| Jersey 1000 Bulls | 47 | 2,608 |

